# Development and validation of a pharmacogenomics reporting workflow based on the Illumina Global Screening Array chip

**DOI:** 10.1101/2023.11.24.568510

**Authors:** Pamela Gan, Muhammad Irfan bin Hajis, Mazaya Yumna, Jessline Haruman, Husnul Khotimah Matoha, Dian Tri Wahyudi, Santha Silalahi, Dwi Rizky Oktariani, Fitria Dela, Tazkia Annisa, Tessalonika Damaris Ayu Pitaloka, Priscilla Klaresza Adhiwijaya, Rizqi Yanuar Pauzi, Robby Hertanto, Meutia Ayuputeri Kumaheri, Levana Sani, Astrid Irwanto, Ariel Pradipta, Kamonlawan Chomchopbun, Mar Gonzalez Porta

## Abstract

**Background:** Microarrays are a well-established and widely adopted technology capable of interrogating hundreds of thousands of loci across the human genome. Combined with imputation to cover common variants not included in the chip design, they offer a cost-effective solution for large-scale genetic studies. Beyond research applications, this technology can be applied for testing pharmacogenomics, nutrigenetics, and complex disease risk prediction.

However, establishing clinical reporting workflows requires a thorough evaluation of the assay’s performance, which is achieved through validation studies. In this study, we performed pre-clinical validation of a genetic testing workflow based on the Illumina Global Screening Array for 25 pharmacogenomic-related genes.

**Methods:** To evaluate the accuracy of our workflow, we conducted multiple pre-clinical validation studies. Here, we present the results of accuracy and precision assessments, involving a total of 73 cell lines. These assessments encompass reference materials from the Genome-In-A-Bottle (GIAB), the Genetic Testing Reference Material Coordination Program (GeT-RM) projects, as well as additional samples from the 1000 Genomes project (1KGP). We conducted an accuracy assessment of genotype calls for target loci in each indication against established truth sets.

**Results:** In our per-sample analysis, we observed a mean analytical sensitivity of 99.39% and specificity 99.98%. We further assessed the accuracy of star-allele calls by relying on established diplotypes in the GeT-RM catalogue or calls made based on 1KGP genotyping. On average, we detected a diplotype concordance rate of 96.47% across 14 pharmacogenomic-related genes with star allele-calls. Lastly, we evaluated the reproducibility of our findings across replicates and observed 99.48% diplotype and 100 % phenotype inter-run concordance.

**Conclusion:** Our comprehensive validation study demonstrates the robustness and reliability of the developed workflow, supporting its readiness for further development for applied testing.

## 1 Introduction

Pharmacogenomics (PGx) is a specialized field of medicine that explores the interplay between an individual’s genetic makeup and their response to medications (Pirmohamed, 2023). It investigates how genetic variants influence drug metabolism, efficacy, and safety, aiding in understanding and predicting individual responses to specific drugs. One prominent example of its application is found in the study of the cytochrome P450 family 2 (CYP2), an extensively researched and well-understood enzyme family responsible for metabolizing approximately 25% of available drugs (Goh et al., 2017). For example, individuals with loss-of-function alleles in CYP2C19 exhibit reduced activation of the prodrug clopidogrel, while those with extra copies of CYP2D6 genes may experience adverse effects from standard doses of codeine (Scott et al., 2013; Crews et al., 2014). By identifying and interpreting these genetic variations, PGx facilitates the development of personalized therapeutic strategies aimed at enhancing drug efficacy and minimizing adverse drug reactions (ADRs).

Studies have shown that up to 70% of ADRs have strong genetic associations (Chan et al., 2016), and the financial burden of trial-and-error prescriptions is estimated to be immense, amounting to USD 30 billion (Sultana et al., 2013). To date, consortia such as the Clinical Pharmacogenetics Implementation Consortium (CPIC) and the Dutch Pharmacogenetics Working Group (DPWG) have published genotype-based guidelines for over a hundred gene-drug pairs, providing a robust and evidence-backed framework to facilitate the integration of PGx into everyday clinical practice (Relling and Klein, 2011; Bank et al., 2018; Relling et al., 2020; Abdullah-Koolmees et al., 2021). Remarkably, it is estimated that over 90% of the population carries at least one actionable pharmacogenomic variant, indicating the vast potential of PGx testing in guiding drug therapy and reducing the risk of ADRs (Dunnenberger et al., 2015; Pirmohamed, 2023). Thus, when contemplating a broad implementation of PGx testing, it is important to select a technology that is both widely accessible and cost-effective. Additionally, as outcomes of the tests can be used to guide therapeutic decisions, it is important to establish the analytical and clinical validity of the results before implementing them into patient care. For many molecular tests, assessing the analytical performance is relatively simple, as they yield binary outcomes like positive or negative results. However, PGx markers present a spectrum of genotypes which, when combined, can lead to diverse phenotypes. For example, CYP enzyme metabolizer phenotypes can range from “poor metabolizers” to “ultra-rapid metabolizers”. Therefore, careful consideration needs to be taken when designing validation studies to endorse the use of PGx tests (Huebner et al., 2023).

In this study, we describe the development and validation of a clinical reporting workflow for pharmacogenomics testing. This workflow uses the Illumina GSA chip, a widely available and cost-effective genetic testing solution, to report on 503 distinct variants across 25 PGx genes associated with 303 clinically actionable drugs. To our knowledge, this is the first comprehensive validation of the Illumina GSA v3 chip for pre-emptive pharmacogenomics testing.

## 2 Materials and Methods

### 2.1 Samples and genotyping

The 3 Genome-In-A-Bottle (GIAB) DNA samples were obtained from the National Institute of Standards and Technology (NIST). In addition, a total of 70 DNA samples were purchased from the Coriell Institute for Medical Research, (https://corielle.org). These samples were selected to cover an array of ethnicities and pharmacogenes (Supplementary Table 2).

Infinium Global Screening Array-24 v3.0 BeadChips were processed according to the standard Infinium High-throughput Screening (HTS) protocol according to manufacturer’s instructions. Signal intensities were converted into idat files using an iScan® machine (Illumina Inc.).

For CNV calling, B-allele frequency (BAF) and log-likelihood (LRR) data were generated using GenomeStudio v2.0.5 (Illumina Inc.) with a custom cluster file created according to the manufacturer’s instructions. Array-based sample genders were also estimated in this step.

### 2.2 Data processing

#### 2.2.1 Quality control

For genotyping, idat files were converted to gtc format based on human genome build GRCh38.p13 using the iaap cli (v1.1.0) followed by conversion to VCF using gtc_to_vcf.py (v1.2.1)(https://github.com/Illumina/GTCtoVCF).

Chip-wide autosomal call rates were calculated using plink2 (v alpha3.7) and only samples with greater than 0.98 call rate and whose array-based genders matched to the expected genders were processed. Sites with >90% missing calls and with less than 0.5% minor allele frequency were removed prior to phasing with Eagle2 (Loh et al., 2016) and imputation with minimac4 (Das et al., 2016) using the 1KGP Phase 3 (Byrska-Bishop et al., 2022) data as a reference panel.

#### 2.2.2 CNV Calling

CNV calls were made with PennCNV (Wang et al., 2007) using a custom pfb file and GC-content model generated using scripts from PennCNV. CNV calls for samples with LRR standard deviation greater than 0.2, BAF drift greater than 0.01 and a wave factor greater than 0.05 after GC model adjustment are not considered further. CNV calls supported by less than 10 probes or covering less than 250 bp were removed using the filter_cnv.pl script from PennCNV.

Reference CNV calls were obtained based on GeT-RM and PacBio calls (Chen et al., 2021). In addition, reference calls for WT copy number, whole gene duplications and deletions were obtained from a 1KGP WGS analyzed dataset generated in a previous study from Lee et al. (2021) (Lee et al., 2022). When a duplication is detected, the CYP2D6 allele that is duplicated cannot be assigned based on the available information and is reported as copy number≥3.

#### 2.2.3 Diplotype Calling

Diplotype calls were made using a combination of PharmCAT (v2.2.3) (Sangkuhl et al., 2020) and in-house custom scripts. When an ambiguous call is made, the call is resolved where possible based on the frequency in the population to which the sample belongs.

#### 2.2.4 Metabolizer Profiles

In general, for genes with diplotype calls, metabolizer profiles were assigned based on diplotype calls referring to a curated set of gene, diplotype and phenotype calls obtained from PharmGKB. For samples with CYP2D6 duplications, metabolizer profiles were assigned after calculation of copy number (Crews et al., 2014).

### 2.3 Assessment of Accuracy and Precision

Genotyping accuracy was assessed using GIAB samples (HG001, HG002, HG005) and 1KGP samples. For 1KGP samples, genotype calls from the 30x 1000 Genome Phase 3 Reanalysis with DRAGEN 3.7.6 accessed from https://registry.opendata.aws/ilmn-dragen-1kgp on November 3, 2023 were used as the truth set. For GIAB samples, DeepVariant genotype calls from UltimaGenomics accessed on June 7, 2023 (Updated standard reference Genome-in-a-Bottle (GIAB) samples HG001-HG007, n.d.) were used as the truth set. Variant calls were classified as true positive, true negative, false positive and false negative as previously described (Kishikawa et al., 2019). Discordant calls in CYP2D6, which is known to have many structural variants, and G6PD, on the X chromosome, were adjusted manually, if supported by external information (expected copy numbers or gender).

The following metrics were calculated as such:

- Callability: The percentage of successfully genotyped loci out of all considered genotypes.
- Genotype concordance: The percentage of genotyped sites with a correct call.
- Analytical sensitivity: The percentage of variant sites correctly identified.
- Analytical specificity: The percentage of non-variant sites correctly identified.
- Precision: The percentage of variants correctly genotyped relative to the number of reported variants.
- No-call rate: Percentage of missing genotypes out of all considered genotypes.

Per-site concordances were calculated using the same definitions for all 503 variants on a per-site basis out of 65 samples per site.

For diplotype concordance, concordance was defined as the percentage of samples with a correct call out of samples with a reference call. For UGT1A1, CYP2D6 and SLCO1B1 genes, diplotype calls that differed between the reference dataset calls and our pipeline were still considered concordant if appropriate based on the pipeline’s reportable range. Specifically, *1 and *2 for CYP2D6 were evaluated as equivalent as the key variant for *2 is not directly genotyped, in the absence of an imputed call, will default to *1. For UGT1A1, *60 was considered equivalent to *1 and *80 was considered a proxy for *28 and *37. For CYP2C19, *38, which is the reference allele reported in the absence of any mutation, was considered equivalent to *1. Diplotypes were considered concordant as long as there were GeT-RM results with the same genotype.

Confidence intervals for point estimates of the above metrics were estimated using the Wilson score interval (Wilson, 1927; Newcombe and Altman, 2011). Confidence intervals for means of the above metrics were estimated by bootstrapping (n=10,000).

#### 2.3.1 Allele frequencies

Alleles frequencies were obtained from the Phase 3 1000 Genomes dataset described above. Samples were subsetted by population and allele frequencies were calculated using bcftools (v1.9) +fill-tags. In addition, population level aggregated allele frequencies per-site were obtained from the Tohoku Medical Megabank project (The Tohoku Medical Megabank Project: Design and Mission, 2016).

## 3 Results

### 3.1 PGx reporting workflow and validation study design

We have developed a pharmacogenomics reporting workflow centered around the Illumina GSA v3 chip, as depicted in **Figure 1**. The patient journey begins with a pre-test counselling session, during which eligible participants provide informed consent, submit buccal samples, and complete an indication-specific survey detailing their current medications. The buccal swab sample is then sent to a clinically accredited laboratory for DNA extraction and genotyping. An in-house bioinformatics analysis pipeline is used to convert raw experimental data from IDAT format into VCF files. Following direct genotyping, imputation is employed to characterize loci not covered by the microarray chip. This step is followed by diplotype determination for target PGx genes. During the final analysis step, diplotyping results are interpreted into metabolizer profiles, and the results are enriched with actionable recommendations drawn from established PGx guidelines. Finally, a PDF report is generated and provided to the patient during a post-test consultation.

**Figure 1:**
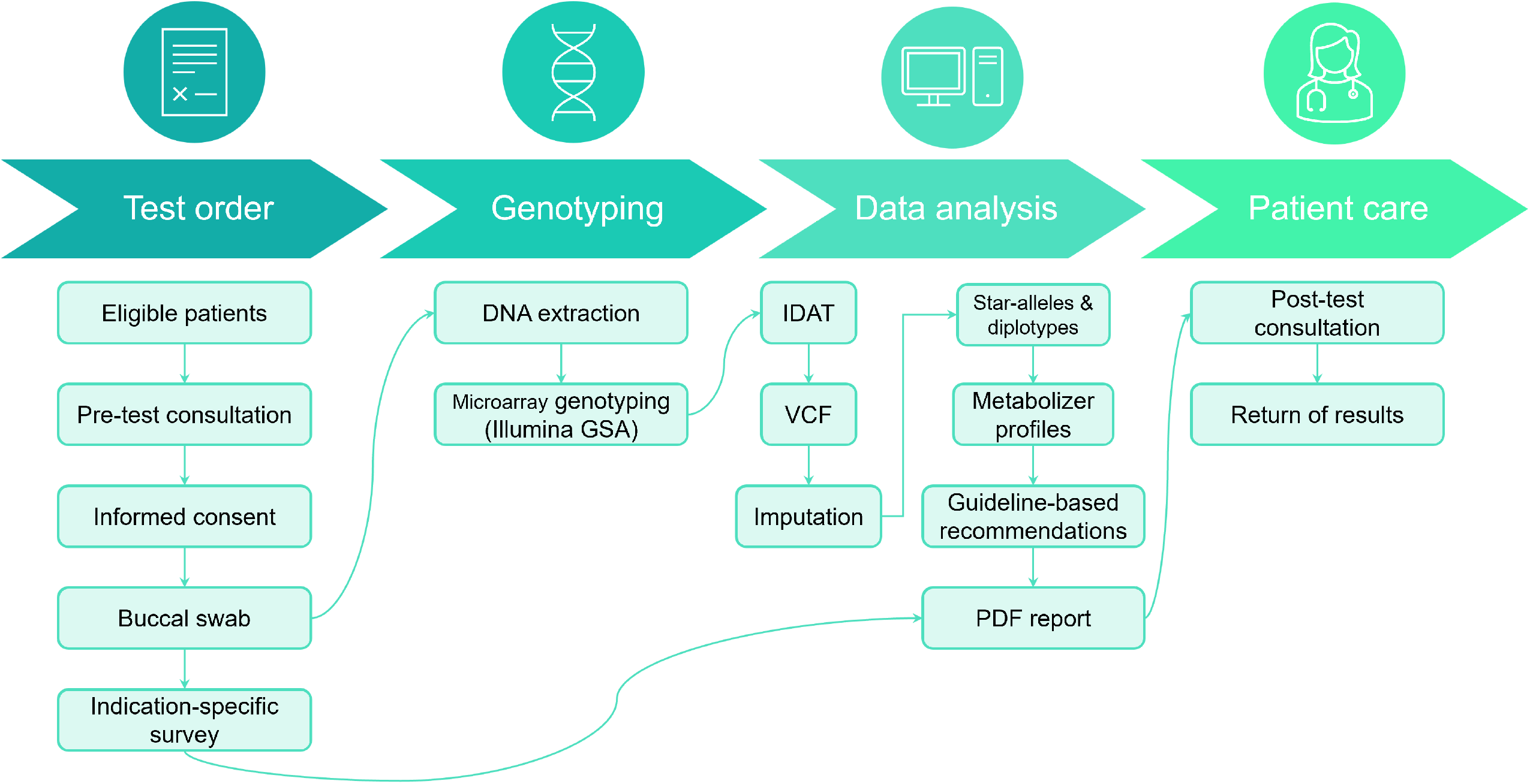
End-to-end PGx testing workflow. During the pre-test consultation, participants provide informed consent, complete a PGx survey, and submit a DNA sample from a buccal swab. This sample undergoes DNA extraction and array genotyping, followed by bioinformatic analysis to characterize variants in selected PGx genes. Genotype calls are subsequently interpreted into metabolizer profiles and annotated with actionable recommendations from published guidelines. Finally, results are compiled into a PDF report, which is discussed with the patient during a post-test consultation.

Our reporting workflow encompasses a total of 25 pharmacogenes and 503 distinct variants corresponding to 429 haplotypes (**Supplementary Table 1**). These genes have been selected to include the Very Important Pharmacogenes listed in PharmGKB which overlap with markers in the Illumina GSA chip (N=21 out of 35 Tier 1 VIPs)(Whirl-Carrillo et al., 2012). To validate the accuracy of the test results, we designed a comprehensive validation study using well-established reference materials, selected to represent a broad range of PGx outcomes (**Figure 2** and **Supplementary Table 2**). Specifically, we utilized 3 GIAB cell lines, 45 cell-lines from the United States Centers for Disease Control and Prevention (CDC) Genetic Testing Reference Material (GeT-RM) Coordination Program (Pratt et al., 2010), including 37 samples that are also part of the 1000 Genomes Project (1KGP) (Byrska-Bishop et al., 2022), and an additional 26 cell lines from the 1KGP to conduct six distinct experiments, which included assessments of accuracy (per-sample, per-site, CYP2D6 CNV calling and star allele concordance) as well as intra-and inter-run reproducibility. Importantly, among the 429 haplotypes included in the reportable range of our test, 84 could be directly tested under this experimental design.

**Figure 2:**
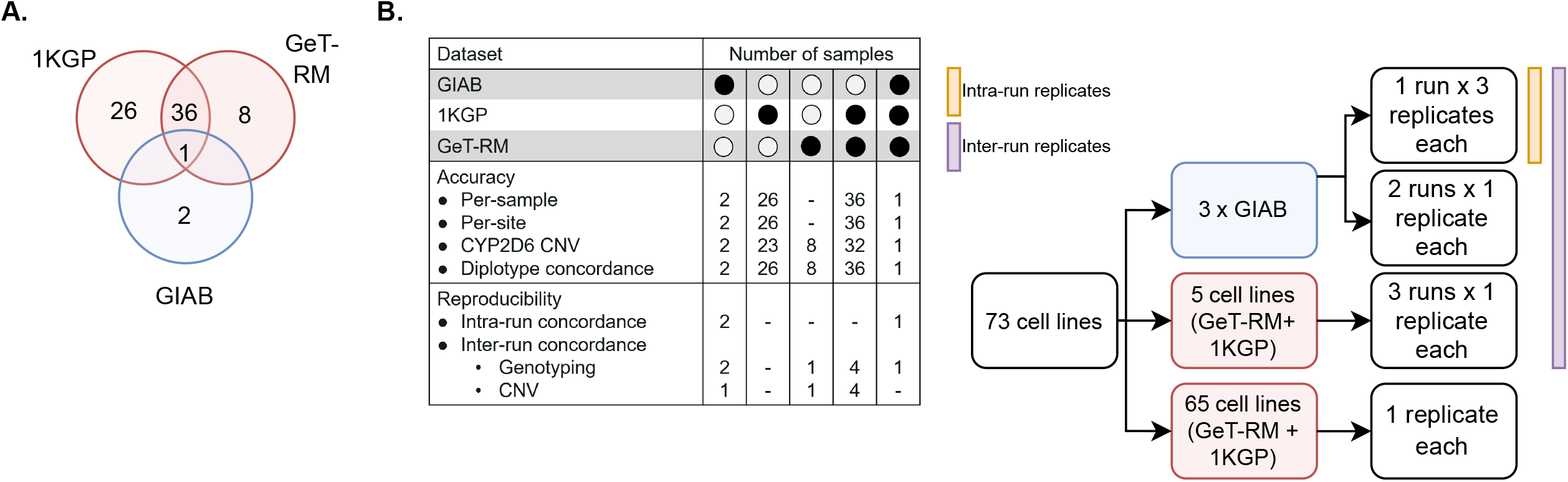
Validation study design. A. DNA from a total of 73 unique reference cell lines were genotyped on the GSA chip to assess genotyping accuracy. Samples were selected due to the availability of well-characterized reference genotype calls (1KGP, GIAB) or reference calls for important PGx genes that have been validated experimentally by multiple labs (GeT-RM). B. Breakdown of samples by experiment. GIAB samples were ran in triplicate in a 3:1:1 design to enable measurement of inter- and intra-run reproducibility. Selected GeT-RM samples with known copy number variations (CNV) were also run across 3 runs to assess inter-run reproducibility of CNV calling

### 3.2 Accuracy assessment of genotype calls

First, we aimed to assess the accuracy of our genotyping and imputation workflow by evaluating our ability to obtain correct genotype calls at individual PGx loci. We conducted two complementary analyses: one centered on per-site assessments and the other on per-sample evaluations.

In the per-site analysis, we evaluated variant calling performance for 503 PGx loci in the 65 samples of the validation set with reference genotype calls. Out of 503 sites, 278 loci had a call in the 1KGP Phase 3 reference set. Of these, 114 sites (41.00 %) could be evaluated with a true positive with the genotyping validation set. For the remaining sites, only 2 (rs114096998, rs34223104) are present in the 1KGP dataset with an allele frequency of higher than 1% and 15 (rs35350960, rs55951658, rs2306282, rs72559747, rs71581941, rs186364861, rs1135835, rs1135833, rs72549352, rs567606867, rs118203758, rs72554664, rs72554665, rs137852342 and rs137852327) have an allele frequency of higher than 1% in at least one of the East Asian populations represented in the dataset (CHS, CDX, KHV, CHB and JPT). Further, only 18 of the unassessed SNPs had a minor allele frequency of greater than 0.1% in the Tommo 54K dataset consisting of 54,300 Japanese individuals, indicating that the majority of unevaluated sites are present at low frequencies in Asian populations.

Of the 114 sites that could be evaluated, only 4 (rs1135840, rs1058164, rs1065852 and rs3093105) had a concordance of less than 95%. Among these, 3 (rs1135840, rs1058164 and rs1065852) were associated with CYP2D6 and, upon closer examination, discrepancies could be attributed to differences in query versus reference dosages. For these sites, the discordant samples were known to harbor structural variants of CYP2D6, and the dosages reported in the reference calls were not consistent with the expected diplotypes. For example, all 3 were reported to be homozygous alternate (dosage=2) in the 1KGP dataset for HG01190 and NA18861, which are known to have only a single copy of the CYP2D6 gene each, while the microarray results reported the expected dosage of 1. This led to the calls being labelled as false negatives, due to the discordance with the truth data. Further, rs1058164 for NA19207, which has a duplicated copy of CYP2D6 *2, was reported as heterozygous according to the reference VCF, whereas the microarray results reported the expected dosage of 2 for the same SNP, thus resulting in misclassification as a false positive. After adjustment, the results indicated that 99.80% of sites (502/503) exhibited 95% concordance with the expected calls (**Figure 3**), thus demonstrating a high level of per-site accuracy in our workflow.

**Figure 3:**
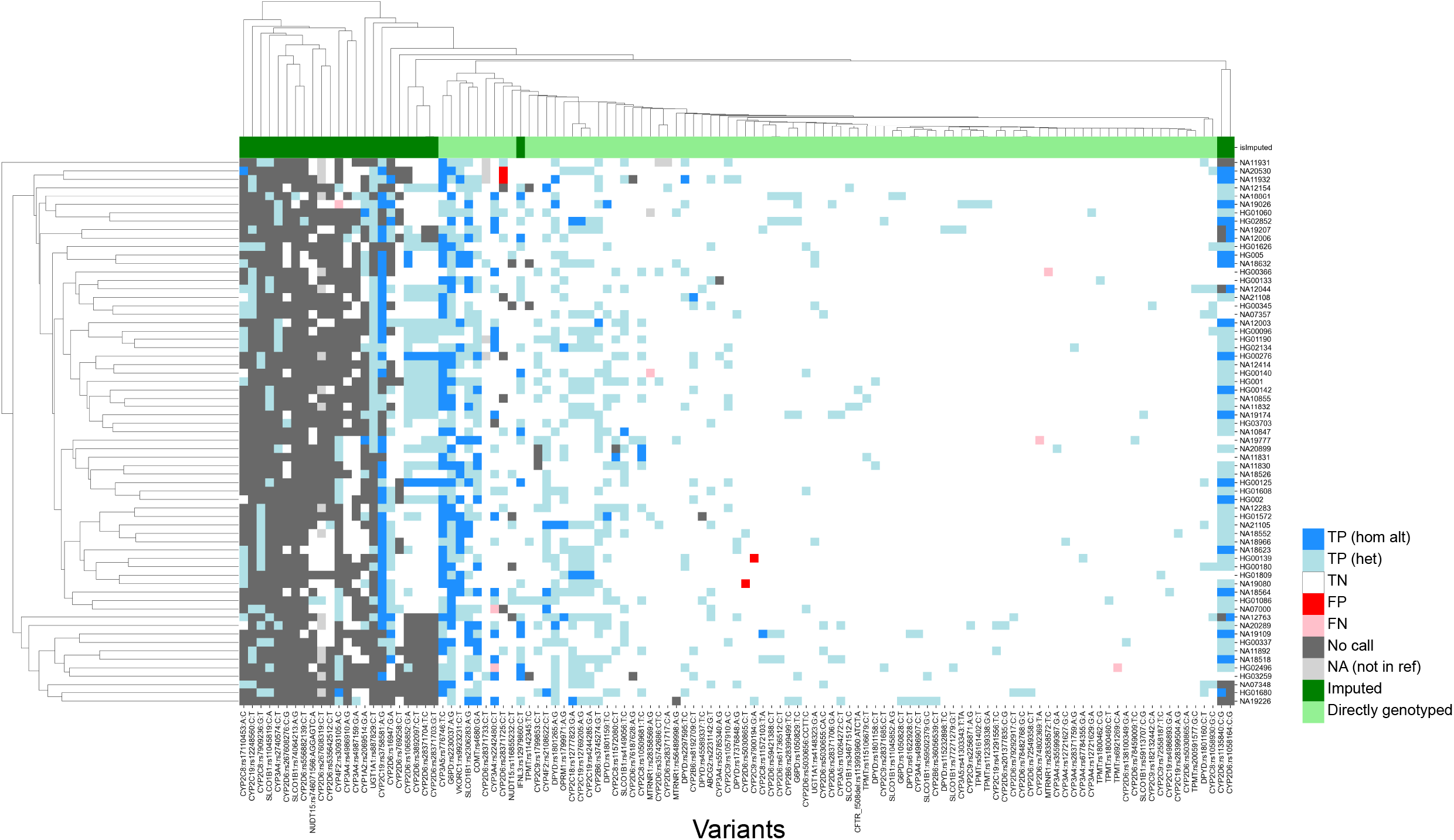
Genotyping concordance against 65 accuracy controls (per-site analysis). Heatmap showing concordance of 114 variants with true positives. 1KGP and GIAB samples were utilized as accuracy controls. Imputed sites have a lower callability compared to sites that are directly genotyped. TP (hom alt): True positive homozygous alternate, TN(het): true positive heterozygous, TN: True negative, FP: false positive, FN: false negative, NA: not in reference

After manual review of discordant calls, our results revealed consistently high analytical sensitivity and specificity across all tested samples, with means of 99.39% [95%CI=91.67-100.00] and 99.98 [95%CI=99.79-100.00], respectively (**Table 1**). Altogether, these findings demonstrate the applicability of our workflow in accurately determining genotypes for small variants at PGx loci.

**Table 1:** Per-sample accuracy assessment (SNPs and INDELs). Mean sample callability, genotype concordance, analytical sensitivity, analytical specificity, precision and no-call rates of 65 samples, focusing on 503 loci, with 95 % confidence intervals estimated by bootstrapping.

| Metric | Mean (%) |
| --- | --- |
| Callability | 96.08 [94.11, 97.45] |
| Genotype concordance | 99.96 [99.79, 100.00] |
| Analytical sensitivity | 99.39 [91.67, 100.00] |
| Analytical specificity | 99.98 [99.79, 100.00] |
| Precision | 99.54 [93.75, 100.00] |
| No-call rate | 3.92 [2.55, 5.89] |

Additionally, we conducted an extensive evaluation of the CYP2D6 gene, which encodes the Cytochrome P450 2D6 enzyme responsible for metabolizing approximately 25% of clinically used drugs. This gene exhibits significant genetic diversity among individuals, including structural variants and complex events like hybrid rearrangements (Zhou et al., 2017; Gaedigk et al., 2018). The presence of two pseudogenes in the human genome further complicates genotyping efforts. In this study, our primary objective was to assess the capability of our assay to accurately genotype complex variants within the CYP2D6 gene. To achieve this, we included 22 cell lines with challenging CYP2D6 haplotypes into our experimental design, encompassing whole gene deletions, duplications, and complex events such as hybrid rearrangements and co-occurring deletions and duplications. Based on this dataset, we estimate the analytical sensitivity and specificity for CNVs in CYP2D6 gene to be 60.00% [95%CI=35.75-80.18] and 100.00% [95% CI=89.85-100.00], respectively (**Table 2**). To troubleshoot the observed decrease in analytical sensitivity, we divided the performance assessment by structural variant type, and identified that the drop in performance was primarily driven by hybrid tandem duplications. After excluding this specific event type from our evaluation, analytical sensitivity and specificity improved to 90.91% [95%CI=62.26-98.38] and 100.00% [95%CI=90.82-100.00], respectively. Consequently, we decided to exclude hybrid rearrangements for CYP2D6 from the reportable range of our assay.

**Table 2:** CYP2D6 structural variation detection accuracy assessment. Concordance, sensitivity, specificity and precision metrics are shown along with the 95% confidence intervals. The assay had a callability of 78.79% [95%CI=67.49-86.92] and a no-call rate of 21.21% [95%CI=13.08-32.51] across the 66 cell lines included in the analysis.

| Structural variation class | Number in reference set | Genotype concordance (%) | Analytical sensitivity (%) | Analytical specificity (%) | Precision (%) |
| --- | --- | --- | --- | --- | --- |
| All | 22 | 88.46<br>[77.03, 94.60] | 60.00<br>[35.75, 80.18] | 100.00<br>[90.59, 100.00] | 100.00<br>[70.09, 100.00] |
| Duplication + Deletion | 17 | 98.08<br>[89.88, 99.66] | 90.91<br>[62.26, 98.38] | 100.00<br>[91.43, 100.00] | 100.00<br>[72.25, 100.00] |
| Duplication (xN) | 9 | 98.08<br>[89.88, 99.66] | 85.71<br>[48.69, 97.43] | 100.00<br>[92.13, 100.00] | 100.00<br>[60.97, 100.00] |
| Deletion (*5) | 8 | 100.00<br>[93.12, 100.00] | 100.00<br>[51.01, 100.00] | 100.00<br>[92.59, 100.00] | 100.00<br>[51.01, 100.00] |
| Hybrid tandem duplication | 8 | 88.46<br>[77.03, 94.60] | 0.00<br>[0.00, 39.03] | 100.00<br>[92.29, 100.00] | N/A |

### 3.3 Concordance of PGx star alleles

Next, we proceeded to evaluate diplotype calling accuracy in 14 out of 25 pharmacogenes that are reported as star-alleles in our test (**Supplementary Table 1**). An inherent challenge of this analysis lies in the extensive number of star-alleles associated with each pharmacogene, best exemplified by CYP2D6, which includes over 100 known star-alleles in PharmGKB (Zhou et al., 2017; Gaedigk et al., 2018). As a result, finding reference materials to cover each of the alleles during validation is a challenging task, and some alleles (e.g., population-specific ones or novel additions) may not even be present in the current reference material resources. To address this challenge, we selected samples from the GeT-RM database to cover as many samples as possible with the highest diversity of star alleles for the pharmacogenes in our test, prioritizing frequent haplotypes. We curated a final validation set of 73 samples, covering star-alleles for 84 out of 429 possible haplotypes that can be identified by the 503 sites in our test, and achieving a coverage rate of 35.40%. Among the genes considered, UGT1A1 displayed the highest coverage level, while G6PD had the lowest coverage (3.39%), aligning with known variation and lack of reference data in GeT-RM, where only 2 out of the 186 non-reference haplotypes are represented, and where the majority of “haplotypes” are single SNP calls (McDonagh et al., 2012).

Having identified an appropriate sample set that maximizes star-allele representation, we proceeded to genotype each sample using our PGx workflow and subsequently evaluated the concordance of predicted diplotype calls against their respective truth sets. Two sources of truth sets were utilized: GeT-RM calls were used as the truth set for CYP2B6, CYP2C19, CYP2C8, CYP2C9, CYP2D6, CYP3A4, CYP3A5, CYP4F2, TPMT and UGT1A1. For DPYD, G6PD, NUDT15 and SLCO1B1, truth sets were generated based on 1KGP NGS VCFs run on PharmCAT. For NUDT15 and G6PD, this was due to the limited number of samples with GeT-RM calls among the validation set, and for DPYD and SLCO1B1, the 1KGP dataset was used as a reference due to updates in the haplotype definitions since the GeT-RM studies were carried out. Per-gene concordances were calculated using only samples that had calls with either the GeT-RM or 1KGP reference truth sets (**Table 3**).

**Table 3:** Concordance of PGx star alleles. Diplotype and phenotype callability and concordance for 14 genes with haplotype calls. For UGT1A1, genotype concordance was adjusted by assuming *60 is equivalent to *1 and that *80 is equivalent to *27 and *37. For CYP4F2, the phenotype concordance is assessed based on if *3 is present or absent since *3 is used for the drug recommendation.

| Gene | Size of reference call set | Diplotype callability (%) | Diplotype concordance (%) | Phenotype callability (%) | Phenotype concordance (%) |
| --- | --- | --- | --- | --- | --- |
| CYP2B6 | 27 | 88.89<br>[71.94, 96.15] | 95.83<br>[79.76, 99.26] | 88.89<br>[71.94, 96.15] | 95.83<br>[79.76, 99.26] |
| CYP2C19 | 27 | 96.30<br>[81.72, 99.34] | 92.31<br>[75.86, 97.86] | 96.30<br>[81.72, 99.34] | 92.31<br>[75.86, 97.86] |
| CYP2C8 | 27 | 100.00<br>[87.54, 100.00] | 100.00<br>[87.54, 100.00] | NA<br>[NA, NA] | NA<br>[NA, NA] |
| CYP2C9 | 27 | 92.59<br>[76.63, 97.94] | 100.00<br>[86.68, 100.00] | 92.59<br>[76.63, 97.94] | 100.00<br>[86.68, 100.00] |
| CYP2D6 | 40 | 80.00<br>[65.24, 89.50] | 87.50<br>[71.93, 95.03] | 90.00<br>[76.95, 96.04] | 97.22<br>[85.83, 99.51] |
| CYP3A4 | 27 | 100.00<br>[87.54, 100.00] | 100.00<br>[87.54, 100.00] | 100.00<br>[87.54, 100.00] | 100.00<br>[87.54, 100.00] |
| CYP3A5 | 27 | 100.00<br>[87.54, 100.00] | 100.00<br>[87.54, 100.00] | 100.00<br>[87.54, 100.00] | 100.00<br>[87.54, 100.00] |
| CYP4F2 | 25 | 100.00<br>[86.68, 100.00] | 92.00<br>[75.03, 97.78] | 100.00<br>[86.68, 100.00] | 100.00<br>[86.68, 100.00] |
| DPYD | 43 | 86.05<br>[72.74, 93.44] | 100.00<br>[90.59, 100.00] | 90.70<br>[78.40, 96.32] | 100.00<br>[91.03, 100.00] |
| NUDT15 | 64 | 100.00<br>[94.34, 100.00] | 100.00<br>[94.34, 100.00] | 100.00<br>[94.34, 100.00] | 100.00<br>[94.34, 100.00] |
| SLCO1B1 | 63 | 96.83<br>[89.14, 99.13] | 93.44<br>[84.32, 97.42] | 98.41<br>[91.54, 99.72] | 100.00<br>[94.17, 100.00] |
| TPMT | 31 | 100.00<br>[88.97, 100.00] | 93.55<br>[79.28, 98.21] | 100.00<br>[88.97, 100.00] | 93.55<br>[79.28, 98.21] |
| UGT1A1 | 25 | 100.00<br>[86.68, 100.00] | 96.00<br>[80.46, 99.29] | 100.00<br>[86.68, 100.00] | 96.00<br>[80.46, 99.29] |
| G6PD | 63 | 100.00<br>[94.25, 100.00] | 100.00<br>[94.25, 100.00] | 100.00<br>[94.25, 100.00] | 100.00<br>[94.25, 100.00] |

The results of this analysis revealed an average diplotype concordance of 96.47% across the 14 genes assessed (**Table 3**). Notably, certain genes, including NUDT15, DPYD, G6PD, UGT1A1, CYP3A5, CYP3A4, CYP2C8, and CYP2C9, displayed a high degree of concordance consistent with the relatively low number of variants under consideration (average of 3 diplotypes), in part due to other genotypes being relatively rare.

In contrast, the CYP2D6 gene displayed the lowest diplotype concordance (87.50%), which was expected due to the aforementioned challenges. Specifically, fusion or hybrid haplotypes, such as *10+*36, *68+*4, were identified as *10 or *4, respectively, which aligns with the lowest performance in detecting CNVs previously discussed. In addition, rarer star alleles such as *15, *82, *17 and *56 were either not identified or were reported as wild type (the absence of a reportable mutation). Further, for SLCO1B1, *37/*37 was incorrectly called as *14/*37 in 5 samples due to the inability to resolve the correct diplotype based on the expected frequency in the population.

Importantly, when interpreting diplotypes into metabolizer profiles, phenotype concordance improved across all genes, with an average observed value of 98.07% (**Table 3**). The same improvement was also detected for CYP2D6 (97.22% phenotype concordance), despite the diplotype call differences due to hybrid/tandem duplications described above. This is most likely due to the most common CYP2D6 hybrid tandem duplications being associated with star alleles with similar allele functions. For example, *10+*36, which is the fusion of a no-function *36 allele and a decreased function *10 allele, is itself a decreased function allele, which is functionally the same as the *10 allele detectable by the pipeline. Another example is *68+*4 (no function allele), which is functionally the same as *4 only, that is reportable by the pipeline. For SLCO1B1, *14/*37 and *37/*37 both evaluated as normal function and overall, the phenotype concordance was 100.00%. The lowest phenotype concordance was CYP2C19 (92.31%).

### 3.4 Precision (reproducibility) study

Finally, we conducted a precision study to evaluate the reproducibility of our workflow, considering both intra-run and inter-run consistency at both the genotype and diplotype levels.

To assess intra-run precision of genotype calls, we utilized three GIAB samples (HG001, HG002, and HG005), each ran in triplicate, achieving 100% concordance across all replicates (**Supplementary Table 3**). In the inter-run evaluation of genotype calls, we expanded the dataset to include five additional GeT-RM samples, also ran in triplicate, consistently observing 100.00% concordance across all runs (**Supplementary Table 3**). These additional GeT-RM samples were deliberately selected because they harbor known duplications and deletions in CYP2D6. Further analysis of intra-run performance for CYP2D6 diplotype calling confirmed 100.00% diplotype calling concordance across all runs (**Supplementary Table 4**). Lastly, by expanding the analysis to include all reported genes, we identified an average of 100.00% intra-run concordance and 99.50% inter-run concordance (**Supplementary Table 5**). The slight reduction in inter-run genotype concordance was influenced by differences in variants called for MT-RNR1 between runs of NA19226. However, phenotype concordance was 100.00% between all samples, including for MT-RNR1 of NA19226 where all samples were assigned a Normal Risk based on the results.

## 4 Discussion

Pharmacogenomics (PGx) is revolutionizing personalized medicine by providing insights into individual drug responses based on genetics. This approach has the potential to significantly improve treatment outcomes, reduce adverse drug reactions, and ultimately lower treatment costs (Pirmohamed, 2023). It is estimated that over 90% of the population carries at least one actionable pharmacogenomic variant (Dunnenberger et al., 2015; Pirmohamed, 2023). Furthermore, PGx information is already actionable: the DPWG and CPIC consortia have to date curated dosing guidelines based on genetics for over 140 drugs (Bank et al., 2018; Abdullah-Koolmees et al., 2021). As such, with the increasing incorporation of PGx information into patient care, there emerges a pressing need for the development and validation of routine PGx tests. Our study introduces a pre-emptive PGx reporting workflow that utilizes the widely available Illumina GSA chip, covering 503 variants across 25 pharmacogenes, and including 21 out of the 35 Tier 1 Very Important Pharmacogenes listed in PharmGKB (those with markers in the Illumina GSA chip).

To assess the accuracy and reliability of our test, we designed a comprehensive validation study, incorporating a selection of well-established reference materials from the GIAB (Genome in a bottle—a human DNA standard, 2015) and GeT-RM consortia (Pratt et al., 2010, 2016, 2022; Gaedigk et al., 2022). Additionally, we included cell lines from the 1000 Genomes Project (Byrska-Bishop et al., 2022) to address loci not covered by the aforementioned resources, but that were part of the reportable range of our test. In total, our study comprised 73 unique samples, strategically chosen to maximize the representation of star-alleles across the 25 genes covered by our test.

We analyzed the results from this sample set to establish the analytical performance of the assay, including per-sample and per-site accuracy, CNV calling performance in CYP2D6, star allele concordance, and as intra- and inter-run reproducibility. From these studies, and focusing on the target 503 PGx loci, we determined the assay’s mean analytical sensitivity to be 99.39% [95%CI=91.67-100.00] and the analytical specificity to be 99.98% [95%CI=99.79-100.00]. To complement these performance metrics, and identify systematic sources of error at each locus, we also conducted a more detailed per-site analysis, evaluating each target site across all samples in the validation set. We observed that 99.80% of loci (502/503) consistently demonstrated concordance with the expected calls across samples. Next, following the interpretation of genotype calls into star-alleles and diplotype calls, we continued our assessment by comparing diplotype results to those in the truth set. On average, for the 14 genes with haplotypes, the diplotype concordance was 96.47%. Notably, the evaluation of a subset of samples in replicate settings yielded consistent results, with an average 99.48% inter-run and 100% intra-run concordance rates. Overall, our workflow was demonstrated to exhibit both accuracy and precision. When compared to other array-based PGx assays, our test exhibited performance levels closely aligned with the reported literature standards for accuracy (ranging from 93% to 100%) and precision (ranging from 97% to 100%)(Hartshorne et al., 2013; Martins et al., 2013; Borobia et al., 2018; Collins et al., 2019; Tang et al., 2021; Kanji et al., 2023).

In addition, our validation results highlight the inherent complexities associated with genotyping specific PGx genes, notably CYP2D6. The genetic diversity of this gene, along with structural variants, hybrid rearrangements, and pseudogenes, makes it challenging to genotype. In our assessment of SV calling performance for CYP2D6, we noted that no signature of a duplication was detected except for one instance of a non-identical whole gene duplication (NA18526, *1/*36x2+*10). In general, such hybrid genes are detected only over specific exon/introns depending on whether the tandem duplication occurs over the 3’ or the 5’ end of the gene. Based on the current settings, the minimum size of a reported CNV is 250 bp, which may be too small to identify most of such duplications. Potentially, reducing the minimum CNV size may allow better detection of such variations, however, it is expected that a higher number of false positives may also be reported. Additionally, diplotype concordance for SV-containing star-alleles in CYP2D6 was lower than that of the other evaluated diplotypes (91.30% if ignoring tandem hybrid samples vs. 87.50% for all samples). These observations led us to exclude this specific variant type from the reportable range of the assay. While our findings align with existing research on the complexities of genotyping certain PGx genes, they also underscore the limitations of microarrays as the definitive platform for PGx reporting. Emerging sequencing technologies, particularly those based on long-read sequencing, are poised to bridge this performance gap and standardize reportable loci, thus minimizing discrepancies across tests (van der Lee et al., 2020). However, such methods remain comparatively more expensive alternatives, and for now are typically reserved for situations where cost-efficiency is not the primary consideration. Our primary objective in this study was to develop a pharmacogenomics test with the potential for broad adoption. To achieve this, we selected the Illumina GSA platform due to its widespread availability and cost-effectiveness. Furthermore, the platform’s capacity to provide genome-wide data opens the door to various additional applications.

Altogether, the notion of pre-emptive PGx testing seamlessly integrating into healthcare frameworks is no longer a distant vision but an impending reality. Numerous research endeavors, including randomized controlled trials, have gathered evidence supporting the customization of drug therapy based on pharmacogenetic testing targeting specific drug-gene interactions to improve patient outcomes (Mallal et al., 2008; Pirmohamed et al., 2013; Coenen et al., 2015; Henricks et al., 2018; Claassens et al., 2019). Additionally, several studies have reported significant reductions in hospital admissions, emergency department visits, and overall healthcare expenditures, indicating the potential cost-effectiveness of genetics-informed treatment approaches (Brixner et al., 2016; Finkelstein et al., 2016; Elliott et al., 2017). Notably, the PREPARE study, conducted across seven European countries with diverse healthcare settings and encompassing a cohort of 6,944 patients, evaluated the impact of genotype-guided prescriptions using a 12-gene pharmacogenetic panel (Swen et al., 2023). This prospective real-world implementation study revealed a 30% reduction in clinically relevant adverse drug reactions when employing a panel-based pharmacogenetic testing strategy.

However, the universal integration of PGx into the standard healthcare landscape is not without its challenges. These encompass a range of practical considerations that extend beyond the accuracy of the underlying genetic tests (Pirmohamed, 2023). Among them, the notable lack of awareness and training among healthcare professionals often leads to suboptimal utilization of pharmacogenomic testing. This knowledge gap extends beyond healthcare practitioners to include patients, who frequently remain uninformed about the potential benefits and accessibility of such tests. Operational challenges are also present and entail the need for streamlined processes for test ordering, result interpretation, and the immediate availability of results to healthcare professionals during patient care. Furthermore, the financial aspect should not be underestimated, particularly in countries lacking uniform insurance coverage, where the cost implications of pharmacogenomic testing can be prohibitive, even with cost-efficient solutions like microarrays. Lastly, navigating the ethical and privacy dilemmas associated with issues such as informed consent, data confidentiality, and the potential for genetic discrimination adds further layers of complexity. Addressing these intricate challenges requires a collaborative effort involving researchers, healthcare experts, policymakers, educators, and various stakeholders. Based on our research, which leverages an easily accessible microarray chip, we maintain an optimistic outlook that our solution, alongside others, will serve as a catalyst for the broader adoption of PGx testing.

## Abbreviations

1KGP: 1000 Genomes Project
ADR: adverse drug reactions
BAF: B-allele frequency
CDC: United States Centers for Disease Control and Prevention
CNV: copy number variations
GIAB: Genome In a Bottle
GSA: global screening array
GeT-RM: Genetic Testing Reference Material
PGx: pharmacogenomics
CPIC: Clinical Pharmacogenetics Implementation Consortium
DPWG: Dutch Pharmacogenetics Working Group
MAF: minor allele frequency
SNP: single nucleotide polymorphism
CHS: Southern Han Chinese
CDX: Chinese Dai in Xishuangbanna, China
KHV: Kinh in Vietnam
CHB: Han Chinese in Beijing
JPT: Japanese people in Tokyo
LRR: log likelihood ratio
CYP: cytochrome P450
MT-RNR1: Mitochondrially Encoded 12S RRNA
NUDT15: Nudix Hydrolase 15
DPYD: Dihydropyrimidine Dehydrogenase
G6PD: Glucose-6-Phosphate Dehydrogenase
UGT1A1: UDP Glucuronosyltransferase Family 1 Member A1
SLCO1B1: Solute Carrier Organic Anion Transporter Family Member 1B1
TPMT: Thiopurine S-methyltransferase

## Ethics statement

Not applicable

## Conflict of Interest

PG, MIH, MY, JH, LS, AI, KC and MGP are employees of NalaGenetics. HKM, DTW, SS, DRO, FD, TA, TDAP, RYP, RH, MAK and AP are employees of Genomik Solidaritas Indonesia.

## Author Contributions

PG, MIH, MY, JH and KC designed, developed and validated the analysis pipeline and reviewed the manuscript. PG analyzed experimental data and was involved in writing the manuscript. AI and LS were involved in funding acquisition and supervised analyses. KC and MGP conceived the study and were involved in writing and reviewing the manuscript. HKM, DTW contributed by investigation, methodology, project management; SS, DR, FD, TA, TDAP, PKA, RYP contributed by investigation, methodology; RH, MAK contributed in terms of funding acquisition, investigation, methodology, resources, supervision, project management; AP contributed by funding acquisition, investigation, methodology, resources, supervision, writing-review.

## Funding

This study was co-funded by NalaGenetics and Genomik Solidaritas Indonesia.

## Acknowledgments

The authors would like to thank internal members at NalaGenetics and Genomik Solidaritas Indonesia for their feedback and inputs.

## Data Availability Statement

The data that supports the findings of this study are available as supplementary materials. Any additional data is available from the corresponding author upon reasonable request.

