## Supplementary Table 1 for "Development and validation of a pharmacogenomics reporting workflow based on the Illumina Global Screening Array chip"

**Supplementary Table 1: Reportable range of the PGx test.**

| Gene | Validated with true positive | Variants/ Haplotypes |
| --- | --- | --- |
| ABCG2 | rs2231142 | rs2231142 |
| CACNA1 | rs1800559, rs772226819 | rs1800559; rs772226819 |
| CFTR | delF508 | delF508; 2789+5G->A; 3272-26A->G; 3849+10kbC->T; 711+3A->G; A1067T; A455E; D110E; D110H; D1152H; D1270N; E56K; E831X; F1052V; F1074L; G1069R; G1349D; G178R; G551D; G551S; K1060T; L206W; P67L; R1070W; R117C; R117H; R347H; R352Q; R74W; S1251N; S1255P; S549R(A>C); S549R(T>G); S945L; S977F |
| COMT | rs4680 | rs4680 |
| CYP1A2 | rs2069514* | rs2069514 |
| CYP2B6 | *2; *6; *18; *20 | *2; *3; (*34, *36); (*6, *7); *8; *11; *12; *13; *15; *17; *18; *19; *20; *21; *22; *23; *26; *27; *28; *35; *37 |
| CYP2C18 | rs12777823 | rs12777823 |
| CYP2C19 | *2; *3; *4; *8; *13; *17; (*1 = *38) | (*1, *38); *2; *3; *4; *5; *6; *7; *8; (*11, *15, *24, *28, *33, *9); *10; *12; *13; *14; *17; *18; *19; *22; *23; *25; *26; *29; *30; *31; *32; *35; *39 |
| CYP2C8 | *2; *3; *4 | *2; *3; *4; *6; *7; *8; *9; *10; *11; *13; *15; *18 |
| CYP2C9 | *2; *3; *8; *9; *61 | *2; *3; *4; *5; *6; *8; *9; *10; *11; *12; *13; *14; *15; *16; *17; *18; *19; *20; *21; *22; *24; *26; *27; *29; *30; *31; *32; *33; *35; *36; *37; *38; *39; *40; *42; *43; *44; *45; *46; *47; *48; *49; *50; *51; *52; *53; *54; *55; *56; *57; *60; *61; *66; *68; *71 |
| CYP2D6* | *3, *4, *5, *6, *9, *10, *17, *28, *29, *33, *35, *41, *99, xN | (*2, *1); *3; *4; *5; *6; *7; *8; *10; *11; *12; *14; *17; *19; *22; *23; *25; *28; *29; *31; *33; *34; *35; *36; *39; *41; *44; *47; *48; *49; *50; *51; *52; *53; *54; *55; *56; *57; *59; *64; *65; *69; *70; (*168,*71); *73; *81; *82; *83; *88; *91; *94; *95; *99; *102; *103; *104; *107; *109; *111; *114; *115; *117; *119; *121; *132; *141; *142; *143; *150; *153; *160; *164 |
| CYP3A4 | *2; *3; *15; *16 | *2; *3; *4; *5; *8; *9; *10; *11; *12; *13; *14; *15; *16; *18; *21; *22; *23; *24; *26; *37; *38 |
| CYP3A5 | *3; *6; *7 | *3; *6; *7; *9 |
| CYP4F2 | *2; *3 | *2; *3 |
| DPYD | *5; *9A | *2A; *3; *4; *5; *6; *7; *8; *9A; *9B; *10; *11; *12; *13; HapB3; c.1057C>T; c.1108A>G; c.1180C>T; c.1181G>T; c.1218G>A; c.1260T>A; c.1294G>A; c.1314T>G; c.1349C>T; c.1358C>G; c.1403C>A; c.1475C>T; c.1484A>G; c.1519G>A; c.1543G>A; c.1577C>G; c.1615G>A; c.1682G>T; c.1775G>A; c.1777G>A; c.1796T>C; c.1896T>C; c.1906A>C; c.1990G>T; c.2021G>A; c.2161G>A; c.2186C>T; c.2195T>G; c.2279C>T; c.2336C>A; c.2482G>A; c.2582A>G; c.2623A>C; c.2639G>T; c.2656C>T; c.2846A>T; c.2872A>G; c.2915A>G; c.2921A>T; c.2933A>G; c.2977C>T; c.2978T>G; c.3049G>A; c.3061G>C; c.3067C>A; c.313G>A; c.451A>G; c.46C>G; c.496A>G; c.498G>A; c.557A>G; c.601A>C; c.61C>T; c.62G>A; c.632A>G; c.775A>G; c.868A>G; c.929T>C; c.934C>T; c.967G>A |
| G6PD | A; A- 202A_376G | A- 202A_376G; A- 680T_376G; Aachen Loma Linda; Ananindeua Hechi Viangchan, Jammu; Bajo Maumere Seattle, Lodi, Modena, Ferrara II, Athens-like; Japan, Shinagawa Kawasaki; Malaga Santa Maria; Mediterranean, Dallas, Panama, Sassari, Cagliari, Birmingham; Puerto Limon; Santiago de Cuba, Morioka; Taipei, Chinese-3; Crispim Salerno Pyrgos Vanua Lava; Alhambra; Andalus; Aveiro; Bangkok Noi Canton, Taiwan-Hakka, Gifu-like, Agrigento-like Cosenza; Beverly Hills, Genova, Iwate, Niigata, Yamaguchi; Cairo; Chatham; Chinese-5; Cincinnati; Cincinnati Minnesota, Marion, Gastonia, LeJeune; Coimbra Shunde Vancouver; Dagua; Gaohe; Gond; Guadalajara Mt Sinai; Harilaou; Hermoupolis Honiara Union,Maewo, Chinese-2, Kalo; Ierapetra; Ilesha; Iowa, Walter Reed, Springfield; Kaiping, Anant, Dhon, Sapporo-like, Wosera; Kalyan-Kerala, Jamnaga, Rohini; Lagosanto; Mahidol; Metaponto; Montpellier; Murcia Oristano; Namouru; Nankang; Nanning; Nashville, Anaheim, Portici; Neapolis; Nilgiri Santiago; Orissa; Pawnee; Riverside; Sao Borja; Serres; Shenzen; Sibari; Sierra Leone; Sinnai; Split; Surabaya; Tomah; Vancouver; Yunan |
| IFNL3 | rs12979860 | rs12979860 |
| MTRNR1 | rs56489998, rs28358569 | rs56489998; rs28358569; rs200887992; rs28358572; rs267606617 |
| NUDT15 | *3 | *2; *3; *4; *5 |
| OPRM1 | rs1799971 | rs1799971 |
| RYR1 |  | rs193922747; rs193922748; rs118192161; rs193922753; rs1801086; rs1801086; rs121918592; rs193922764; rs118192116; rs118192162; rs111888148; rs193922768; rs193922770; rs118192172; rs193922772; rs118192175; rs118192176; rs118192177; rs118192177; rs112563513; rs193922802; rs193922803; rs193922807; rs193922809; rs121918593; rs28933396; rs118192124; rs193922816; rs118192122; rs28933397; rs121918594; rs118192178; rs118192178; rs193922818; rs193922832; rs193922843; rs118192167; rs121918595; rs193922876; rs193922878; rs118192168; rs63749869; rs118192170 |
| SLCO1B1 | *5; *14; *15; *20; *27; *37; *41; *43 | *2; *4; *3; *5; *6; *7; *8; *9; *10; *11; *12; *13; *14; *15; *16; *19; *20; *23; *24; *25; *26; *27; *28; *29; *30; *31; *32; *33; *34; *37; *40; *41; *43; *44; *45; *46; *47 |
| TPMT | *2; *3C; *8; *21; *32; *33 | *2; *3A; *3B; *3C; *4; *5; *7; *8; *9; *10; *11; *13; *14; *15; *17; *18; *19; *20; *21; *23; *24; *25; *26; *27; *28; *29; *30; *31; *32; *33; *34; *35; *36; *37; *38; *39; *41 |
| UGT1A1 | *6; *80 | *6; *80 |
| VKORC1 | rs9923231 | rs9923231 |
