## Supplementary Table 2 for "Development and validation of a pharmacogenomics reporting workflow based on the Illumina Global Screening Array chip"

**Supplementary Table 2: List of samples used in the validation study.**

| Sample ID | Gender | | Ethnicity | Description | GIAB | 1KGP | GeT-RM |
| --- | --- | --- | --- | --- | --- | --- | --- |
| HG001 (NA12878) | F | | Utah/Mormon | Genotyping accuracy and reproducibility control, diplotype control | 1 | 1 | 1 |
| HG002 (NA24385, GM26105, GM27730) | M | | Ashkenazim Jewish | Genotyping accuracy and reproducibility control | 1 | 0 | 0 |
| HG005 (NA24631, GM26107) | M | | Chinese | Genotyping accuracy and reproducibility control | 1 | 0 | 0 |
| HG00096 | M | British from the United Kingdom | | Genotyping accuracy control | 0 | 1 | 0 |
| HG00125 | F | British from the United Kingdom | | Genotyping accuracy control | 0 | 1 | 0 |
| HG00133 | F | British from the United Kingdom | | Genotyping and diplotype accuracy control | 0 | 1 | 1 |
| HG00139 | M | British from the United Kingdom | | Genotyping and diplotype accuracy control | 0 | 1 | 0 |
| HG00140 | M | British from the United Kingdom | | Genotyping accuracy control | 0 | 1 | 0 |
| HG00142 | M | British from the United Kingdom | | Genotyping accuracy control | 0 | 1 | 0 |
| HG00180 | F | Finnish in Finland | | Genotyping accuracy control | 0 | 1 | 0 |
| HG00276 | M | Finnish in Finland | | Genotyping accuracy control, CYP2D6 CNV reproducibility control | 0 | 1 | 1 |
| HG00337 | F | Finnish in Finland | | Genotyping and diplotype accuracy control | 0 | 1 | 1 |
| HG00345 | M | Finnish in Finland | | Genotyping accuracy control | 0 | 1 | 0 |
| HG00366 | M | Finnish in Finland | | Genotyping accuracy control | 0 | 1 | 0 |
| HG01060 | M | Puerto Rican in Puerto Rico | | Genotyping and diplotype accuracy control | 0 | 1 | 0 |
| HG01086 | F | Puerto Rican in Puerto Rico | | Genotyping and diplotype accuracy control | 0 | 1 | 1 |
| HG01190 | M | Puerto Rican in Puerto Rico | | Genotyping and diplotype accuracy control | 0 | 1 | 1 |
| HG01572 | F | Peruvian in Lima, Peru | | Genotyping accuracy control | 0 | 1 | 0 |
| HG01608 | M | Iberian populations in Spain | | Genotyping accuracy control | 0 | 1 | 0 |
| HG01626 | F | Iberian populations in Spain | | Genotyping accuracy control | 0 | 1 | 0 |
| HG01680 | M | Iberian populations in Spain | | Genotyping and diplotype accuracy control | 0 | 1 | 1 |
| HG01809 | F | Chinese Dai in Xishuangbanna, China | | Genotyping and diplotype accuracy control | 0 | 1 | 1 |
| HG02134 | M | Kinh in Ho Chi Minh City, Vietnam | | Genotyping and diplotype accuracy control | 0 | 1 | 0 |
| HG02496 | M | African Caribbean in Barbados | | Genotyping and diplotype accuracy control | 0 | 1 | 1 |
| HG02852 | F | Gambian in Western Division - Mandinka | | Genotyping and diplotype accuracy control | 0 | 1 | 1 |
| HG03259 | F | Gambian in Western Division - Mandinka | | Genotyping and diplotype accuracy control | 0 | 1 | 1 |
| HG03703 | F | Punjabi in Lahore, Pakistan | | Genotyping and diplotype accuracy control | 0 | 1 | 1 |
| NA07000 | F | Utah residents with ancestry from northern and western Europe from the CEPH collection | | Genotyping and diplotype accuracy control | 0 | 1 | 1 |
| NA07348 | F | Utah residents with ancestry from northern and western Europe from the CEPH collection | | Genotyping and diplotype accuracy control | 0 | 1 | 1 |
| NA07357 | M | Utah residents with ancestry from northern and western Europe from the CEPH collection | | Genotyping and diplotype accuracy control | 0 | 1 | 1 |
| NA07439 | F | African-American | | Diplotype calling accuracy control | 0 | 0 | 1 |
| NA10847 | F | Utah residents with ancestry from northern and western Europe from the CEPH collection | | Genotyping and diplotype accuracy control | 0 | 1 | 1 |
| NA10855 | F | Utah residents with ancestry from northern and western Europe from the CEPH collection | | Genotyping and diplotype accuracy control | 0 | 1 | 1 |
| NA11830 | F | Utah residents with ancestry from northern and western Europe from the CEPH collection | | Genotyping accuracy control | 0 | 1 | 0 |
| NA11831 | M | Utah residents with ancestry from northern and western Europe from the CEPH collection | | Genotyping accuracy control | 0 | 1 | 0 |
| NA11832 | F | Utah residents with ancestry from northern and western Europe from the CEPH collection | | Genotyping and diplotype accuracy control | 0 | 1 | 1 |
| NA11892 | F | Utah residents with ancestry from northern and western Europe from the CEPH collection | | Genotyping accuracy control | 0 | 1 | 0 |
| NA11931 | F | Utah residents with ancestry from northern and western Europe from the CEPH collection | | Genotyping accuracy control | 0 | 1 | 0 |
| NA11932 | M | Utah residents with ancestry from northern and western Europe from the CEPH collection | | Genotyping accuracy control | 0 | 1 | 0 |
| NA12003 | M | Utah residents with ancestry from northern and western Europe from the CEPH collection | | Genotyping and diplotype accuracy control | 0 | 1 | 1 |
| NA12006 | F | Utah residents with ancestry from northern and western Europe from the CEPH collection | | Genotyping and diplotype accuracy control | 0 | 1 | 1 |
| NA12044 | F | Utah residents with ancestry from northern and western Europe from the CEPH collection | | Genotyping and diplotype accuracy control | 0 | 1 | 1 |
| NA12154 | M | Utah residents with ancestry from northern and western Europe from the CEPH collection | | Genotyping and diplotype accuracy control | 0 | 1 | 1 |
| NA12283 | F | Utah residents with ancestry from northern and western Europe from the CEPH collection | | Genotyping accuracy control | 0 | 1 | 0 |
| NA12414 | F | Utah residents with ancestry from northern and western Europe from the CEPH collection | | Genotyping accuracy control | 0 | 1 | 0 |
| NA12763 | F | Utah residents with ancestry from northern and western Europe from the CEPH collection | | Genotyping accuracy control | 0 | 1 | 0 |
| NA17169 | F | African-American | | Diplotype calling accuracy control | 0 | 0 | 1 |
| NA17244 | M | Utah residents with ancestry from northern and western Europe from the CEPH collection | | Diplotype calling accuracy and CYP2D6 CNV reproducibility control | 0 | 0 | 1 |
| NA17287 | M | Utah residents with ancestry from northern and western Europe from the CEPH collection | | Diplotype calling accuracy control | 0 | 0 | 1 |
| NA17448 | F | Mexican-American community of Los Angeles | | Diplotype calling accuracy control | 0 | 0 | 1 |
| NA18518 | F | Yoruba in Ibadan, Nigeria | | Genotyping and diplotype accuracy control | 0 | 1 | 1 |
| NA18526 | F | Han Chinese in Beijing, China | | Genotyping and diplotype accuracy control | 0 | 1 | 1 |
| NA18552 | F | Han Chinese in Beijing, China | | Genotyping and diplotype accuracy control | 0 | 1 | 1 |
| NA18564 | F | Han Chinese in Beijing, China | | Genotyping and diplotype accuracy control | 0 | 1 | 1 |
| NA18623 | M | Han Chinese in Beijing, China | | Genotyping accuracy control | 0 | 1 | 0 |
| NA18632 | M | Han Chinese in Beijing, China | | Genotyping and diplotype accuracy control | 0 | 1 | 1 |
| NA18861 | F | Yoruba in Ibadan, Nigeria | | Genotyping and diplotype accuracy control | 0 | 1 | 1 |
| NA18966 | M | Japanese in Tokyo | | Genotyping and diplotype accuracy control | 0 | 1 | 1 |
| NA19026 | M | Luhya in Webuye, Kenya | | Genotyping and diplotype accuracy control | 0 | 1 | 1 |
| NA19080 | F | Japanese in Tokyo | | Genotyping accuracy control | 0 | 1 | 0 |
| NA19109 | F | Yoruba in Ibadan, Nigeria | | Genotyping accuracy control, CYP2D6 CNV reproducibility control | 0 | 1 | 1 |
| NA19174 | M | Yoruba in Ibadan, Nigeria | | Genotyping and diplotype accuracy control | 0 | 1 | 1 |
| NA19178 | M | Yoruba in Ibadan, Nigeria | | Diplotype calling accuracy control | 0 | 0 | 1 |
| NA19207 | M | Yoruba in Ibadan, Nigeria | | Genotyping accuracy control, CYP2D6 CNV reproducibility control | 0 | 1 | 1 |
| NA19226 | M | Yoruba in Ibadan, Nigeria | | Genotyping accuracy control, CYP2D6 CNV reproducibility control | 0 | 1 | 1 |
| NA19777 | M | Mexican-American community of Los Angeles | | Genotyping and diplotype accuracy control | 0 | 1 | 1 |
| NA20289 | F | African-American | | Genotyping and diplotype accuracy control | 0 | 1 | 1 |
| NA20530 | F | Toscani in Italy | | Genotyping accuracy control | 0 | 1 | 0 |
| NA20899 | F | Gujarati Indian | | Genotyping accuracy control | 0 | 1 | 0 |
| NA21105 | M | Gujarati Indian | | Genotyping and diplotype accuracy control | 0 | 1 | 1 |
| NA21108 | F | Gujarati Indian | | Genotyping accuracy control | 0 | 1 | 0 |
| NA23348 | F | Unknown | | Diplotype calling accuracy control | 0 | 0 | 1 |
| NA23877 | M | Unknown | | Diplotype calling accuracy control | 0 | 0 | 1 |
