## Supplementary Table 3 for "Development and validation of a pharmacogenomics reporting workflow based on the Illumina Global Screening Array chip"

**Supplementary Table 3: Intra- and inter-run concordance of genotype calls (SNPs and INDELs).** NA sites are defined as those with no calls in at least 2 samples, which were not assessed for concordance.

| **Sample ID** | **Intra-run precision**  **(N=3 replicates)** | | | **Inter-run precision**  **(N=3 replicates)** | | |
| --- | --- | --- | --- | --- | --- | --- |
|  | **N concordant sites** | **%**  **concordance** | **NA sites** | **N concordant sites** | **% concordance** | **NA sites** |
| HG001 | 480/480 | 100.00 | 23 | 480/480 | 100.00 | 23 |
| HG002 | 482/482 | 100.00 | 21 | 482/482 | 100.00 | 21 |
| HG005 | 484/484 | 100.00 | 19 | 484/484 | 100.00 | 19 |
| HG00276 | - | - | - | 483/483 | 100.00 | 20 |
| NA19226 | - | - | - | 473/473 | 100.00 | 30 |
| NA19109 | - | - | - | 477/477 | 100.00 | 26 |
| NA19207 | - | - | - | 483/483 | 100.00 | 20 |
| NA17244 | - | - | - | 491/491 | 100.00 | 12 |
