## Supplementary Table 4 for "Development and validation of a pharmacogenomics reporting workflow based on the Illumina Global Screening Array chip"

**Supplementary Table 4: Inter-run concordance of genotype calls (SVs in CYP2D6).**

| **Sample** | **CYP2D6 diplotype** | **Expected**  **CYP2D6**  **copy number** | **SV call** | | | **% concordance** |
| --- | --- | --- | --- | --- | --- | --- |
|  |  |  | **Run1** | **Replicate 2** | **Replicate 3** |  |
| HG00276 | *4/*5 | 1 | Del | Del | Del | 100 |
| NA19226 | *2/*2XN | 3 | Dup | Dup | Dup | 100 |
| NA19109 | *2X/*29 | 3 | Dup | No-call | Dup | 100 |
| NA19207 | *2/*10/*XN | 3 | Dup | Dup | Dup | 100 |
| NA17244 | *2/*4/*XN | 4 | Dup | Dup | Dup | 100 |
| HG002 | *2/*4 | 2 | WT | No-call | WT | 100 |
