## Supplementary Table 5 for "Development and validation of a pharmacogenomics reporting workflow based on the Illumina Global Screening Array chip"

**Supplementary Table 5: Intra- and inter-run concordance of all PGx genes reported.**  Diplotypes for CYP2B6, CYP2C19, CYP2C8, CYP2C9, CYP2D6, CYP3A4, CYP3A5, CYP4F2, DPYD, NUDT15, SLCO1B1, TPMT, UGT1A1 and G6PD were assessed across 3 intra-run replicates of GIAB samples, and 3 inter-run replicates of the same GIAB samples plus 5 cell lines. For CYP1A2, COMT, OPRM1, VKORC1, IFNL3, ABCG2, CYP2C18, CFTR, MT-RNR1 and for CACNA1/RYR1 all assessed alleles were compared.

| **Sample ID** | **Intra-run precision**  **(N=3 replicates)** | | **Inter-run precision**  **(N=3 replicates)** | |
| --- | --- | --- | --- | --- |
|  | **N concordant calls** | **%**  **concordance** | **N concordant calls** | **% concordance** |
| HG001 | 24/24 | 100.00 | 24/24 | 100.00 |
| HG002 | 24/24 | 100.00 | 24/24 | 100.00 |
| HG005 | 24/24 | 100.00 | 24/24 | 100.00 |
| HG00276 | - | - | 23/23 | 100.00 |
| NA19226 | - | - | 24/24 | 100.00 |
| NA19109 | - | - | 24/24 | 100.00 |
| NA19207 | - | - | 24/24 | 100.00 |
| NA17244 | - | - | 23/24 | 95.83 |
